# Joint exome and metabolome analysis in individuals with dyslexia: Evidence for associated dysregulations of olfactory perception and autoimmune functions

**DOI:** 10.1101/2024.06.27.600448

**Authors:** Rohit Nandakumar, Xiaojian Shi, Haiwei Gu, Yookyung Kim, Wendy H. Raskind, Beate Peter, Valentin Dinu

## Abstract

Dyslexia is a learning disability that negatively affects reading, writing, and spelling development at the word level in 5%-9% of children. The phenotype is variable and complex, involving several potential cognitive and physical concomitants such as sensory dysregulation and immunodeficiencies. The biological pathogenesis is not well-understood. Toward a better understanding of the biological drivers of dyslexia, we conducted the first joint exome and metabolome investigation in a pilot sample of 30 participants with dyslexia and 13 controls. In this analysis, eight metabolites of interest emerged (pyridoxine, kynurenic acid, citraconic acid, phosphocreatine, hippuric acid, xylitol, 2-deoxyuridine, and acetylcysteine). A metabolite-metabolite interaction analysis identified Krebs cycle intermediates that may be implicated in the development of dyslexia. Gene ontology analysis based on exome variants resulted in several pathways of interest, including the sensory perception of smell (olfactory) and immune system-related responses. In the joint exome and metabolite analysis, the olfactory transduction pathway emerged as the primary pathway of interest. Although the olfactory transduction and Krebs cycle pathways have not previously been described in dyslexia literature, these pathways have been implicated in other neurodevelopmental disorders including autism spectrum disorder and obsessive-compulsive disorder, suggesting the possibility of these pathways playing a role in dyslexia as well. Immune system response pathways, on the other hand, have been implicated in both dyslexia and other neurodevelopmental disorders.

## Introduction

Dyslexia is a learning disability that negatively affects reading, writing, and spelling development at the word level (Lyon et al., 2003) in 5%-9% of children (Francks et al., 2002; Pennington & Bishop, 2009; Peterson et al., 2007). In English speakers, it is often diagnosed using standardized tests of sight word recognition, which requires a stable representation of irregularly spelled word forms (*e.g.*, “would,” “enough”) in memory, and nonword decoding, which requires sequentially sounding out letter sequences in the left-right reading frame based on the regular orthography of English (e.g., “gily,” “chelth”). Some of the most frequently cited associated deficits are in the areas of phonemic awareness (Catts, 1989; Giofrè et al., 2019), attention deficit hyperactivity disorder (ADHD) (Germano et al., 2010; Hendren et al., 2018), difficulty with sequential processing across various task types (Bogaerts et al., 2015; Cowan et al., 2017; Peter, Albert, & Gray, 2020; Peter, Albert, Panagiotides, et al., 2020; Peter et al., 2018), and slowed processing speed (Cardillo et al., 2017; Peter, Matsushita, et al., 2011; Shanahan et al., 2006; Stoodley & Stein, 2006), indicative of phenotypic complexity beyond the deficits in reading and spelling.

Dyslexia is thought to be under genetic influence, but despite decades of research, the mechanisms of genetic disruptions are not yet well understood. Dyslexia appears to be highly genetically heterogeneous, as described in various studies and reviews (Carrion-Castillo et al., 2013; Deriziotis & Fisher, 2017; Gialluisi et al., 2020; Graham & Fisher, 2015; Guerra & Cacabelos, 2019; Mascheretti et al., 2017; Peter, Matsushita, et al., 2011; Rahul & Ponniah, 2021; Scerri & Schulte-Korne, 2010), and as yet an association with the underlying brain functions are only evident for a small number of candidate genes and regions (Carrion-Castillo et al., 2019, 2020; Giraud & Ramus, 2013; Mascheretti et al., 2017). Similar to autism spectrum disorder (ASD), a neurodevelopmental disorder that shares some cognitive impairments and neuroanatomical differences with dyslexia (Stoodley, 2014; Tsermentseli et al., 2008), dyslexia can be caused by rare disruptions of large effect or by multiple common disruptions, each of small effect. The latter type of causality can make gene discovery challenging with traditional methods such as variant filtering. In such cases, tools developed for quantitative trait loci such as genome-wide association studies (GWAS) (Dai et al., 2020) (Doust et al., 2022), gene ontology analysis (Wen et al., 2016), or a combination of these (Anney et al., 2011) may be more effective. Also similar to ASD and other neurodevelopmental conditions, several genes of interest have been classified into functional groups, for example, signal transduction, neuronal migration, cell adhesion, developmental biology (Paracchini et al., 2016). Despite these advances, we lack an understanding of how dyslexia is inherited and how individual genes contribute to specific expression profiles of dyslexia, as mediated by the brain (Black et al., 2017; Xia et al., 2017).

A second area of uncertainty is the role of associated health impairments, such as immunodeficiency (Hugdahl et al., 1990; Tonnessen et al., 1993), sensory dysregulation (“neural noise hypothesis,” neural hyperexcitability) (Hancock et al., 2017), and cerebellar motor dysfunction (Marchand-Krynski et al., 2017; Marien et al., 2014). A comprehensive investigation of these associations in the context of genetic and environmental risk factors has not yet been undertaken. The fact that in the past, studies have either reported or failed to report evidence for certain traits, for example, evidence (Frank & Levinson, 1973; Marien et al., 2014; Nicolson et al., 1995; Peter, Albert, & Gray, 2020; Peter, Albert, Panagiotides, et al., 2020) or no evidence for the cerebellar hypothesis (Ramus et al., 2003; Stoodley & Stein, 2013), and evidence (Degrandi et al., 2021; Hugdahl et al., 1990) or no evidence for immune system disorders (Tonnessen et al., 1993), has led some to interpret the findings as controversial (Macdonald et al., 2017; Scerri & Schulte-Korne, 2010). In contrast, we (Peter, Albert, & Gray, 2020; Peter, Albert, Panagiotides, et al., 2020) and others (Scerri & Schulte-Korne, 2010) propose a spectrum view of dyslexia where certain expressions of dyslexia and associated health impairments represent trait profiles under genetic and environmental influence.

Specifically regarding the immune system, the earliest mention of an association with dyslexia was a study on co-occurrences of left-handedness, immune diseases, and learning disabilities (Geschwind & Behan, 1982). This study gave rise to the Geschwind-Behan-Galaburda hypothesis that posited that prenatal exposure to testosterone influenced brain lateralization, thus mediating handedness, immune system functions, and learning (dis-)abilities (Behan & Geschwind, 1985). This hypothesis was initially met with skepticism, citing a lack of empirical support (Bryden et al., 1994). Indeed, in a sample of 734 Norwegian children, a significant association was found between handedness and dyslexia, and between handedness and immune system disorders, but not between dyslexia and immune disorders (Tonnessen et al., 1993). Conversely, in a sample of 105 children with dyslexia and typical controls, significantly more immune- and autoimmune disorders were found in the dyslexia group, compared to the control group (Hugdahl et al., 1990). In a sample of 74 men with autoimmune thyroid disease (Graves’ disease and Hashimoto’s thyroiditis), higher frequencies of mixed dominance and dyslexia traits were found, compared to controls without autoimmune thyroid disease (Wood & Cooper, 1992). A recent study of autoimmunity in children with dyslexia reported significantly higher levels of antithyroid antibodies, compared to typical peers, supporting the thyroid autoimmunity hypothesis in dyslexia (Degrandi et al., 2021).

As mentioned, sensory dysregulation is an area of interest in dyslexia. When listening to spoken language, the signal processing machinery involves a cascading flow of information through stages of sensory transduction, encoding, storing (sensory, short-term/working, and long-term memory), word recognition, and semantic and syntactic assembly toward extracting the construct of the verbal message (Levelt, 1999, p. 199; Levelt et al., 1999, p. 199; Peter et al., 2018; Stackhouse & Wells, 1997). Here, sensory processing is situated at the start of this cascade. The neural noise hypothesis of dyslexia can be allocated to the sensory encoding process. This type of sensory dysregulation implies neural hyperexcitability and is thought to interfere with the process of recognizing relevant information in a signal as novel, suppressing responses to irrelevant information, and thus extracting the essence of the stimulus for transfer into long-term memory. It has been linked to two genes, *DCDC2* and *KIAA0319,* in an animal model (Che et al., 2014, 2016) and in humans, biochemical associations with cross-modal tasks have been observed (Del Tufo et al., 2018). Two recent studies have investigated the ability of adults with dyslexia to filter out redundant information. In an fMRI study, adults with dyslexia showed sustained high levels of brain activity despite repetitive stimuli, whereas typical controls showed attenuated brain activity in the repetitive condition. These neural adaptation patterns were observed regardless of stimulus type (e.g., written words, images of objects, images of faces) (Perrachione et al., 2016). We showed evidence of diminished neural adaptation in adults with dyslexia using an event related potential (ERP) gating paradigm. Participants listened to pairs of pure tone stimuli presented at 500 ms intervals. The typical controls showed attenuated response amplitudes in response to the second of the two tones, whereas the adults with dyslexia did not, mirroring the diminished attenuation in the fMRI study (Perrachione et al., 2016). The magnitude of the gating effect was associated with the accuracy of performance in a word discrimination task (Peter, McCollum, et al., 2019). Together, these two studies support the hypothesis that individuals with dyslexia may have difficulty filtering out redundant information, extracting the essence of stimuli, and forming stable word representations in long-term memory. If insufficient gating of information leads to sensory overstimulation, this may explain the high rates of comorbidity between dyslexia and ADHD. (Germano et al., 2010; Hancock et al., 2017; Perrachione et al., 2016; Peter, McCollum, et al., 2019)

Studies investigating other aspects of sensory processing such as visual-spatial (Franceschini et al., 2012; Giofrè et al., 2019; Gori & Facoetti, 2014), auditory-speech (Benson et al., 2020; Schulte-Körne et al., 1998), multisensory (Hairston et al., 2005; Mascheretti et al., 2017; Meilleur et al., 2020), or sequential (Bogaerts et al., 2015; Peter, Albert, & Gray, 2020; Peter et al., 2018) processing in individuals with dyslexia suggested the possibility that basic neurobiological processes are disrupted in ways that explain how the cognitive or perceptual precursors to reading are impaired (Perrachione et al., 2016). Sensory processing is a broad term that generally refers to the handling of sensory information by the neural system, including the functions of receptor organs and the peripheral and central nervous systems (Jorquera-Cabrera et al., 2017). It should be noted that relatively little is known about the senses of smell and taste in neurodevelopmental disorders (Reilly et al., 2021).

As mentioned, a small number of candidate regions and genes has been identified (Carrion-Castillo et al., 2013; Gialluisi et al., 2020; Graham & Fisher, 2015; Guerra & Cacabelos, 2019; Mascheretti et al., 2017; Peter, Raskind, et al., 2011; Raskind et al., 2012; Scerri & Schulte-Korne, 2010). However, DNA is not the only biological agent that may influence the phenotype. Metabolomics, a rapidly evolving field focused on measuring many metabolites in parallel in a biological system, may contribute new insights into the biology of dyslexia (Gieger et al., 2008). Metabolomics is a comprehensive approach seeking to understand the regulation of metabolic pathways and networks of a biological system (Kaddurah-Daouk & Krishnan, 2009). Because the metabolome is sensitive to biochemical changes caused by even subtle endogenous/exogenous influences (West et al., 2014), studying it can provide a functional readout of the physiological state of the human body (Gieger et al., 2008) and help identify disorder-specific biomarkers.

To date, few studies have examined neurometabolite concentrations in individuals with dyslexia. Initial work found differences between dyslexic adult males and controls in the left temporo-parietal lobe and right cerebellum, specifically lower choline/N-acetylaspartate (NAA) ratio with decreased bilateral choline and increased NAA in the right and decreased NAA in the left cerebellum (Rae et al., 1998). These findings are in contrast with those of another recent study where dyslexic adult males showed a lower ratio of NAA/choline in the right cerebellar hemisphere together with a higher ratio of choline/creatine in the left cerebellar hemisphere (Laycock et al., 2008). Further, initial work suggested that children with dyslexia showed a larger regional distribution of metabolic activation, characterized by elevated lactate in the left anterior quadrant during a phonological task (Richards et al., 1999). Two studies using proton magnetic resonance spectroscopy (MRS) in preschool children (Lebel et al., 2016) and young adults (Bruno et al., 2013) also measured phonological processing ability and demonstrated that phonological processing ability was positively associated with glutamate, creatine, and inositol concentrations in the anterior cingulate in typical preschool children (Lebel et al., 2016), while the other study demonstrated that in the left angular gyrus choline/creatine ratio is negatively correlated with phonological awareness (Bruno et al., 2013). The positive correlation reported between phonological ability and glutamate suggested by previous research (Bruno et al., 2013) is also in the opposite direction to the finding from the longitudinal sample with children with and without dyslexia (Pugh et al., 2014), showing a negative correlation of choline and glutamate concentration in the occipital cortex with reading performance and phonological ability. The authors suggested that the pattern of elevated glutamate in dyslexia might reflect neuronal hyperexcitability contributing to learning deficits in dyslexia (Pugh et al., 2014). In a subsequent study of children, mentioned above (Del Tufo et al., 2018), performance on a cross-modal word matching task was found to mediate the relationship between increased glutamate and poor reading ability, as well as the relationship between increased choline and poor reading ability.

The authors point out that glutamate is proposed to be an indicator of neural noise. Another recent MRS study reported that children with dyslexia, especially females, showed significant negative correlations between processing speed and myo-inositol and choline concentration suggesting that metabolite changes in the anterior cingulate cortex may be present in children with dyslexia and may hold value as a possible marker for dyslexia (Horowitz-Kraus et al., 2018). Another study that investigated adults and children, both with and without dyslexia, suggested that individuals with dyslexia have significantly lower total N-acetylaspartate (tNAA) than controls in the occipital cortex. Adults with dyslexia compared to children with dyslexia were characterized by higher choline and creatine, higher tNAA in left temporo-parietal, and lower glutamate in the visual cortex (Kossowski et al., 2019).

Although recent studies successfully identified dysregulated metabolites associated with dyslexia, there is still a gap in knowledge as to the significance of metabolite concentrations in brain regions. The inconsistencies in previous studies might be attributed to small sample sizes, different participant age groups (pre-reading children vs. school aged children vs. adults), differences in reference scaling (absolute concentration vs. reference to creatine or NAA) or different brain regions examined (occipital lobe vs. cerebellum vs. left temporoparietal region) (Kossowski et al., 2019). Thus, metabolomics research in dyslexia is only in its nascent stage at present, and there is no straight-forward interpretation of these metabolic results (Ramus et al., 2018). 2

An enhanced understanding of the biological substrates of dyslexia in general aligns with the tenets of Precision Medicine, an approach to understanding human health and disease with a purview that includes biological, environmental, and lifestyle related factors toward improved outcomes using personalized treatment and prevention strategies (Khoury & Dotson, 2021).

Translating knowledge of biological substrates into proactive and personalized intervention is common in medical disorders such as cancer and diabetes, but only emerging in developmental disorders with behavioral manifestations such as disorders of spoken and written language.

We present results of a pilot study that queries the intersection of genetics and metabolomics in dyslexia. Biological domain knowledge, such as biological pathway information or protein-protein interactions (PPI), can supplement statistical and data mining approaches to identify biomarkers associated with disease (Khoury & Dotson, 2021) (Dinu, Zhao, et al., 2007; Subramanian et al., 2005). These approaches are particularly useful in the study of complex disorders, such as neurological disorders, cancer, diabetes, etc., which could be influenced by the interaction of multiple genes and environmental factors. We used methods we developed for incorporating domain knowledge in the analysis of association between genomic variants and disease (Dinu, Zhao, et al., 2007) that we successfully applied in etiologic studies of various disorders (Bradley et al., 2014; Briones & Dinu, 2012; Day et al., 2016; Dinu, Miller, et al., 2007;

Gallitano et al., 2012; Huentelman et al., 2015; Lancaster et al., 2020; Li & Dinu, 2018; Peter, Dinu, et al., 2019; Xiang et al., 2008). Our preliminary findings for dyslexia encourage further investigation of metabolites that are related with the Krebs cycle, olfactory transduction, and immune pathways in larger samples.

## Participants and Methods

### Participants

This study was conducted with the oversight and approval of the Institutional Review Boards at the University of Washington and Arizona State University. Adults gave written consent, parents gave written permission for their minor children to participate, and school-age children gave written assent. The exome and metabolome data were collected at two timepoints separated by five years.

Participants were recruited via local clinics providing services to individuals with learning disabilities, the University of Washington Disability Resources for Students, flyers posted on the University of Washington campus, an announcement on co-author BP’s lab website, and word of mouth. Interested individuals and families contacted co-author BP for an initial phone call to determine eligibility. The original consent form included the option of being recontacted for future studies and all participants in the present study had checked this option.

Inclusionary criteria for dyslexia status required a professional diagnosis and performance below -1 SD on at least one of five tests of written language. These tests were administered at the time of the exome data collection and consisted of the Word Identification and Word Attack subtests from the Woodcock Reading Mastery Tests – Revised (Blackwell, 2001), the Sight Word Reading Efficiency and the Phonemic Decoding Efficiency subtests from the Test of Word Reading Efficiency (TOWRE) (Tarar et al., 2015), and the spelling subtest from the Wechsler Individual Achievement Test – Second Edition (Wechsler, 2005). To be included as typical control required performance on all five tests better than -1 SD. In all participants, presence of any medical or sensory condition that could confound the results (e.g., Trisomy 21, hearing impairment) was excluded. However, other health conditions such as ADHD and immune system deficiency were not ruled out because these were relevant for the hypotheses guiding this study.

Of 48 participants whose exomes and/or metabolomes were analyzed, 36 represented 2- or 3-generational families with dyslexia (average family size = 5.14, range [4, 7]. In addition, 10 unrelated adults with, and 2 without, dyslexia participated. From 4 of the families, 5 participants were removed from this study because their dyslexia status could not be unambiguously determined (they either had a professional diagnosis but performed above the cutoff in all tests of reading and spelling, or they had no diagnosis of dyslexia but performed below the cutoff in one or more tests). From the group of participants that qualified for inclusion, exome sequences were available for 38, metabolic data for 28, and both types of data for 23. Exomes were available for 16 males and 13 females with dyslexia (average age 33.4 years, SD = 21.6) and 3 males and 6 females without dyslexia (average age 44.0 years, SD = 22.8). Metabolic data were available for 11 males and 6 females with dyslexia (average age 33.7 years, SD – 19.9) and 4 males and 5 females without dyslexia (average age 49.0 years, SD = 24.7). The Appendix lists the participants by family constellation, sex, affectation status, ages at exome and metabolome collection, and available specimen type, with annotations for significant self-reported conditions such as difficulties with attention and/or focus, sensory dysregulation, and autoimmune disease; note that one participant with dyslexia had Hashimoto’s disease.

## Metabolomics

### Sample Collection and Preparation

Saliva samples (2 ml) were collected using conical tubes and stored on ice in transit to a -80℃ freezer. Transit duration was less than 2 hours. The following reagents were used: Acetonitrile (ACN), methanol (MeOH), ammonium acetate, and acetic acid, all LC-MS grade, were obtained from Fisher Scientific (Pittsburgh, PA). Ammonium hydroxide was obtained from Sigma-Aldrich (Saint Louis, MO). DI water was obtained in-house by a Water Purification System from EMD Millipore (Billerica, MA). PBS was obtained from GE Healthcare Life Sciences (Logan, UT). The standard compounds corresponding to the measured metabolites were obtained from Sigma-Aldrich and Fisher Scientific.

Frozen saliva samples were thawed overnight under 4°C, and 100 μL of each saliva sample was transferred into a 2 mL Eppendorf vial. In preparation for protein precipitation and metabolite extraction, 500 μL MeOH and 50 μL internal standard solution (containing 1,810.5 μM ^13^C3-lactate and 142 μM ^13^C5-glutamic acid) were added to each sample. The mixture was vortexed for 10 s and stored at -20°C for 30 min, followed by centrifugation at 14,000 RPM for 10 min at 4° C. The supernatants (450 μL) were transferred into a new Eppendorf vial and dried using a CentriVap Concentrator (Labconco, Fort Scott, KS). The dried samples were reconstituted in 150 μL of 40% PBS/60% ACN. The quality-control sample was a pooled sample of all saliva samples.

The detailed targeted LC-MS/MS method was described in a growing number of studies (Carroll et al., 2015, p. 20; Eghlimi et al., 2020; Gu et al., 2015, 2016; Jasbi et al., 2019; Shi et al., 2019). All LC-MS/MS procedures were performed on an Agilent 1290 UPLC-6490 QQQ-MS (Santa Clara, CA) system. Each sample was injected twice, 10 µL for analysis using negative ionization mode and 4 µL for analysis using positive ionization mode. Both chromatographic separations were performed in hydrophilic interaction chromatography (HILIC) mode on a Waters XBridge BEH Amide column (150 x 2.1 mm, 2.5 µm particle size, Waters Corporation, Milford, MA). The flow rate was 0.3 mL/min, auto-sampler temperature was kept at 4 ℃, and the column compartment was set at 40 ℃. The mobile phase was composed of Solvents A (10 mM ammonium acetate, 10 mM ammonium hydroxide in 95% H2O/5% ACN) and B (10 mM ammonium acetate, 10 mM ammonium hydroxide in 95% ACN/5% H2O). After the initial 1 min isocratic elution of 90% B, the percentage of Solvent B decreased to 40% at t=11 min. The composition of Solvent B maintained at 40% for 4 min (t=15 min), and then the percentage of B gradually returned to 90% in preparation for the next injection. Mass spectrometry was performed using an electrospray ionization (ESI) source. Multiple-reaction-monitoring (MRM) mode was used for targeted data acquisition. Agilent Masshunter Workstation software (Santa Clara, CA) was used to control the LC-MS system and to integrate the extracted MRM peaks. *Metabolite Data Analysis* Statistical analysis of the metabolites was performed using MetaboAnalyst (Pang et al., 2021) (Pang et al., 2021). A data file with Patient IDs, metabolite levels of 295 metabolites, and class label was entered into the analysis. Using the “Missing Values” option, features with greater than 50% missing values were removed and remaining missing values were replaced by 1/5 of the minimum positive value for each variable. Further filtering was applied based on interquartile range. A log10 transformation was then performed on the data. Volcano plots, which are a combination of fold change (FC) analysis (horizontal axis) and unpaired t-tests (vertical axis) for each metabolite, were charted, and, following the default settings in MetaboAnalyst, metabolites with a raw p-value threshold of 0.1 and fold change threshold of 2.0 were selected as metabolites of interest.

## Exomes

### Sample Collection and Sequencing Methodology

Saliva samples were collected using Oragene DNA OG-500 kits (DNA Genotek, Ottawa, Canada). DNA was extracted using standard procedures as recommended by the manufacturer. Exome sequencing was performed at the University of Washington Center for Mendelian Genomics. Library construction and exome capture were performed using automation (Perkin-Elmer Janus II) in 96-well plate format. 500 ng of genomic DNA was subjected to a series of shotgun library construction steps, including fragmentation through acoustic sonication (Covaris), end-polishing, and A-tailing, ligation of sequencing adaptors, and PCR amplification with dual 10bp barcodes for multiplexing. Prior to sequencing, the library concentration was determined by fluorometric assay and molecular weight distributions verified on the Agilent Bioanalyzer (consistently 150 ± 15bp). Barcoded exome libraries were pooled using liquid handling robotics prior to loading. Massively parallel sequencing-by-synthesis with fluorescently labeled, reversibly terminating nucleotides was carried out on a NovaSeq sequencer.

The sequencing pipeline was a combined suite of Illumina software and other software packages (i.e., Genome Analysis ToolKit [GATK], Picard, BWA, SAMTools, and in-house custom scripts) and consisted of base calling, alignment, local realignment, duplicate removal, quality recalibration, data merging, variant detection, genotyping, and annotation. Variant detection and genotyping were performed using the HaplotypeCaller (HC) tool from GATK (3.7). Variant data for each sample were formatted (variant call format [VCF]) as “raw” calls that contain individual genotype data for one or multiple samples and flagged using the filtration walker (GATK) to mark sites that were of lower quality/false positives [e.g., low quality scores (Q50), allelic imbalance (ABHet 0.75), long homopolymer runs (HRun> 3), and/or low quality by depth (QD < 5)].

Data QC included an assessment of: (1) total PE75 reads; (2) library complexity; (3) capture efficiency; (4) coverage distribution: 90% at 8X required for completion; (5) capture uniformity; (6) raw error rates; (7) Transition/Transversion ratio (Ti/Tv); (8) distribution of known and novel variants relative to dbSNP (typically < 7%); (9) fingerprint concordance > 99%; (10) sample homozygosity and heterozygosity; and (11) sample contamination validation. Exome completion is defined as having > 90% of the exome target at > 8X coverage and > 80% of the exome target at > 20X coverage. Typically, this requires mean coverage of the target at 50-60X.

Variants were annotated using the SeattleSeq Annotation Server (Ng et al., 2009). This publicly accessible server returns annotations including dbSNP rsID (or whether the coding variant is novel), gene names and accession numbers, predicted functional effect (e.g., splice-site, nonsynonymous, missense, etc.), protein positions and amino-acid changes, PolyPhen predictions, conservation scores (e.g., PhastCons, GERP), ancestral allele, dbSNP allele frequencies, and known clinical associations.

### Exome Analysis

Allelic association analysis was run on PLINK (Purcell et al., 2007), where 38 patients’ exome variants were loaded. An R script was developed to further analyze the unadjusted *p*-value file obtained from the PLINK association analysis. SNPs with a *p*-value under 0.05 were selected. The SNP rs IDs were aligned to HGNC gene symbol and ENSEMBL IDs. Potentially novel SNPs that did not have rs IDs were entered into BioMart by chromosome location to identify genes by Gene Symbol and ENSEMBL IDs. The gene symbols and ENSEMBL IDs for SNPs with or without rsID were concatenated, respectively, and frequency count analysis was performed to identify genes that occurred most frequently. A ranked score was developed based on dividing the number of variants per gene with the size of that gene’s exon-coding sequence. This number, generally very small, was then multiplied by 10,000 to help create ranked scores that are easier to interpret. This score provided a normalized metric of hits per gene. GOrilla (Eden et al., 2009) was then used to identify overexpressed GO terms by entering a ranked list of genes that were identified in the previous step.

### Combined Metabolite and Exome Analysis

The gene list from conducting the exome analysis and the top metabolite list identified with MetaboAnalyst were used as inputs into the Joint Pathway Analysis option in MetaboAnalyst. Only genes that had a ranked score of > 1 were selected, which allowed for the inclusion of 260 high-scoring genes, as MetaboAnalyst does not score/rank genes automatically. We chose a cutoff for the ranked scores to be 1 or greater, as this allows only a small fraction of genes to be selected while still retaining enough genes to conduct analyses. Selected analysis parameters were integration of all pathways using a hypergeometric test with degree centrality and integration of all *p* values.

### Network Analysis

In the network analysis, the compound list inputted is the top metabolite list identified in our statistical analysis. The “Metabolite-Metabolite Interaction Network” option was selected and default settings were used to generate the network analysis plots.

## RESULTS

### Metabolites

Nine metabolites (pyridoxine, kynurenic acid, stachyose hydrate, citraconic acid, phosphocreatine, hippuric acid, xylitol, 2-deoxyuridine, and acetylcysteine) met nominal statistical significance in the metabolite statistical analysis (Table 1, Figure 1, Figure 2). As stachyose hydrate was not recognized by the MetaboAnalyst package in the Joint Pathway Analysis or the Network Analysis modules, this metabolite was removed prior to conducting subsequent analyses. Table 1 lists the metabolites by fold change and statistical significance metrics. Figure 1 is a volcano plot of the metabolites by fold change and significance metrics. Figure 2 shows box plots of the metabolite distributions in the participants with dyslexia, compared to the unaffected participants. Figure 3 is a visualization of the network analysis of the metabolites of interest.

**Figure 1:**
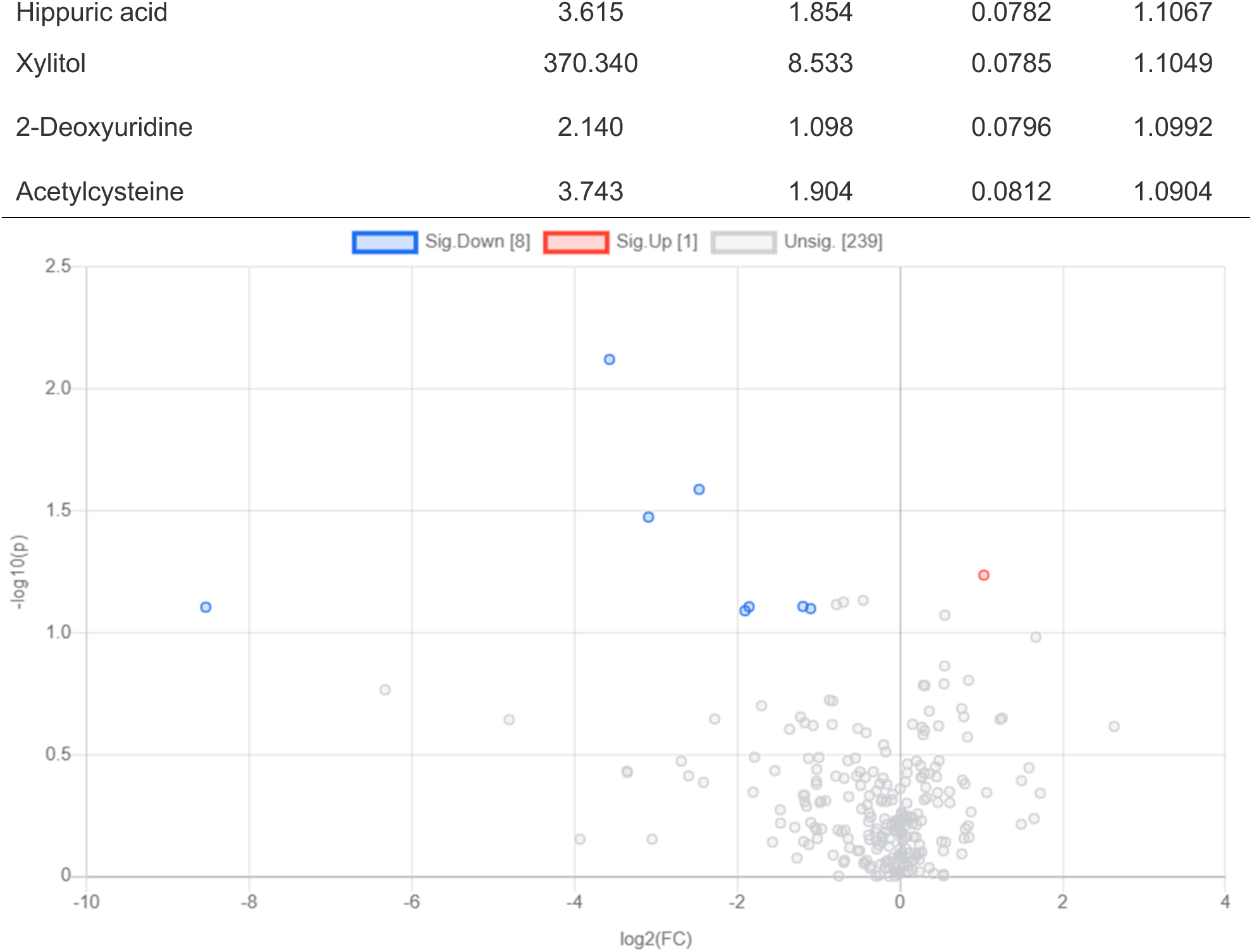
Volcano plot with a fold change threshold of 2.0 and p-value threshold of 0.10. Both the blue and red highlighted dots are metabolites that passed these threshold filters. The blue dots represent metabolites that are significantly lower in metabolite levels in our case (dyslexic) group, while the red dots represent metabolites that are significantly higher in metabolite levels in our case group compared to the control group.

**Figure 2.**
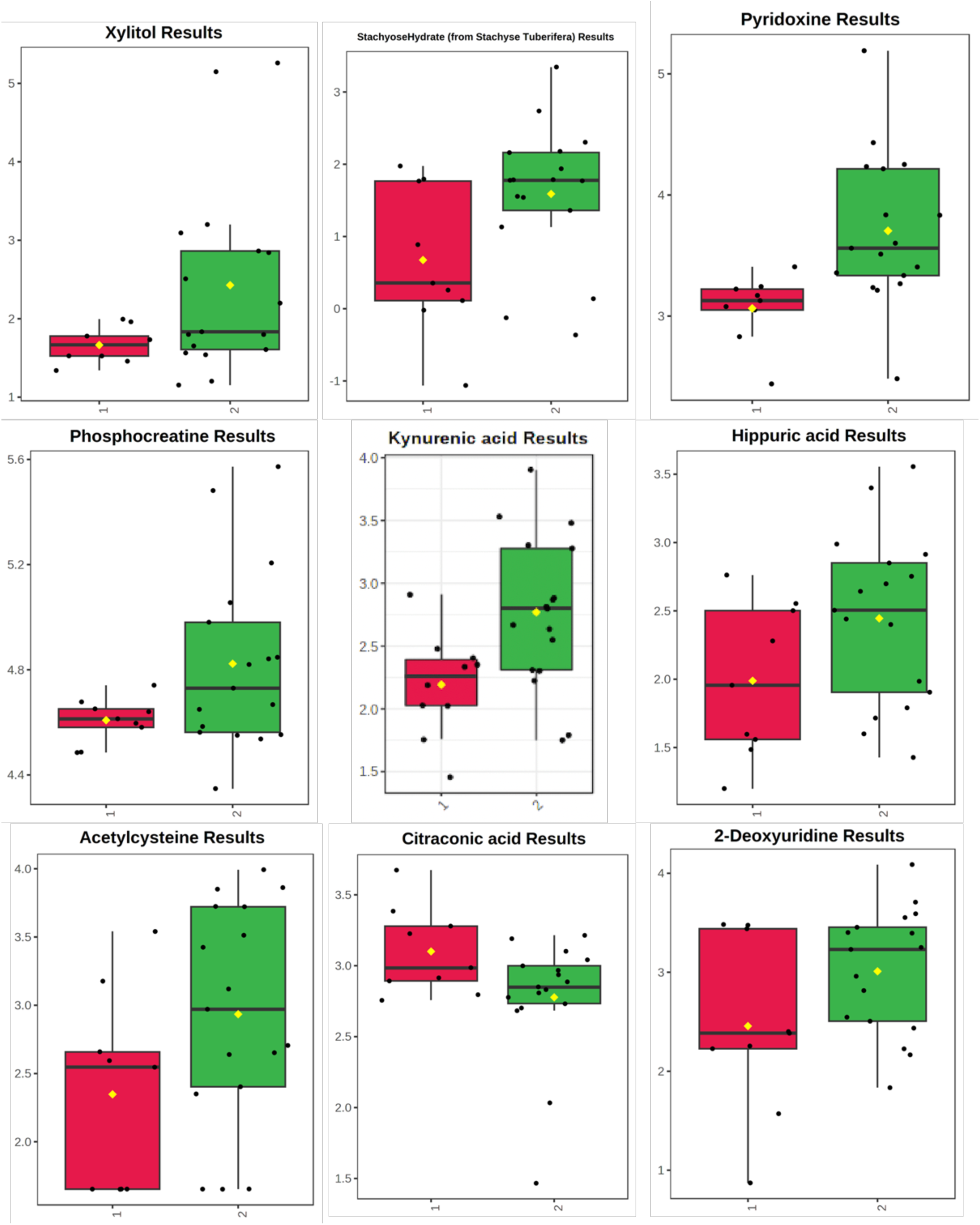
Kynurenic acid, pyridoxine, acetylcysteine, citraconic acid, 2-deoxyuridine, hippuric acid, phosphocreatine, stachyse hydrate, and xylitol distributions for control individuals (red, left, labeled 1) and individuals with dyslexia (green, right, labeled 2). The black dots are the metabolite expression levels of a given metabolite for each individual, with each individual’s metabolite levels plotted as a black dot. Average levels are denoted by yellow dots.

**Figure 3.**
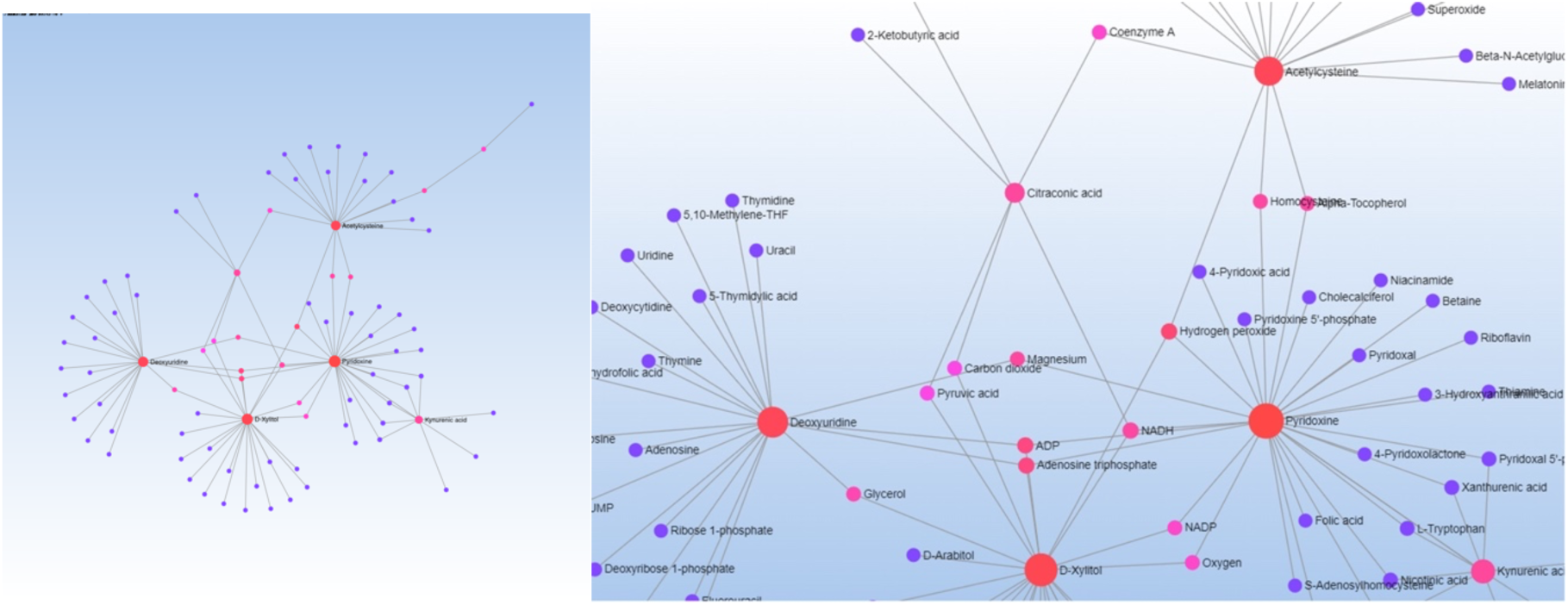
Visualization of the network analysis of the metabolites of interest performed in MetaboAnalyst. (a) Zoomed-in image of metabolite network analysis with a focus on the Krebs intermediates. (b) Metabolite network analysis showing all interactions.

**Table 1.**
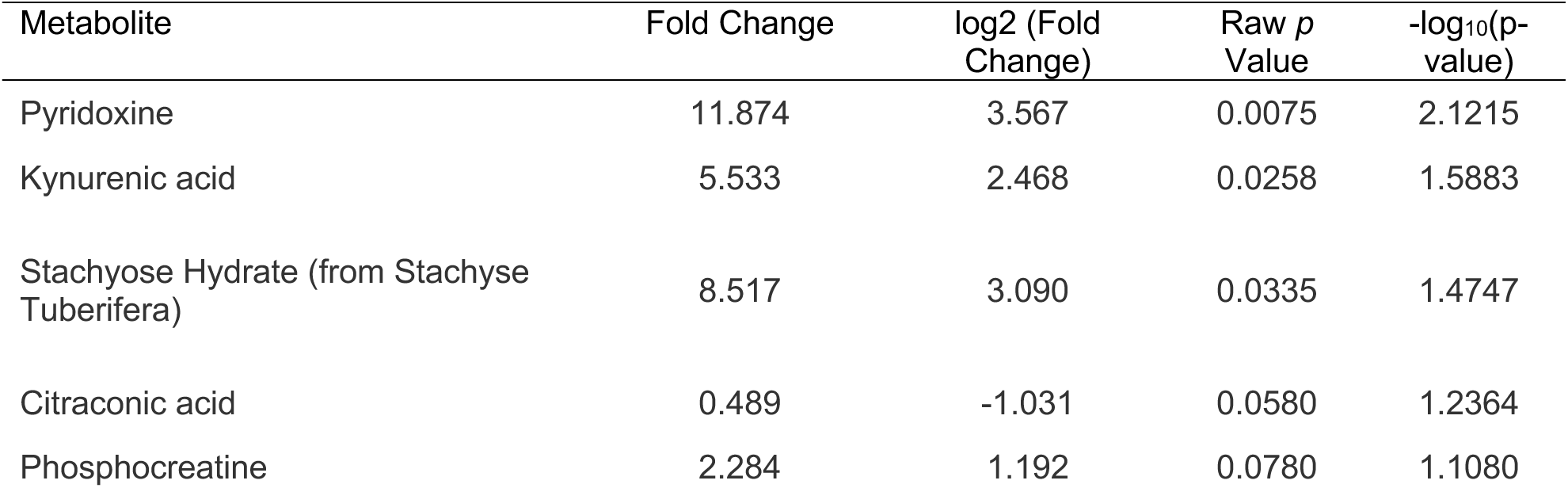
Metabolites with the highest statistical significance, all with raw p-value < 0.10, the log-transformed values, and fold change threshold of > 2.0 or < 0.5.

### Exomes

The association analysis using PLINK and R resulted in 1,813 SNPs below the *p*-value threshold of 0.05. Choosing a more inclusive threshold for the p-value, such as .05, uncorrected for multiple testing, is a common practice when performing pathway analysis to allow the inclusion of multiple signals from the genome. Of these, 181 SNPs did not have rs ids. Pathway analysis with GOrilla resulted in twenty gene ontology (GO) terms with a false discovery rate (FDR) *q*-value < .01. Of these, six involve sensory perception, nine directly relate to the immune system (including antigen processing), and three involve signal transduction. Table 2 lists the GO terms, their descriptions, *p*-values, FDR *q*-values, and enrichment metrics. Figure 4 shows the GO visualization of the genes of interest identified from PLINK and our R analysis.

**Figure 4.**
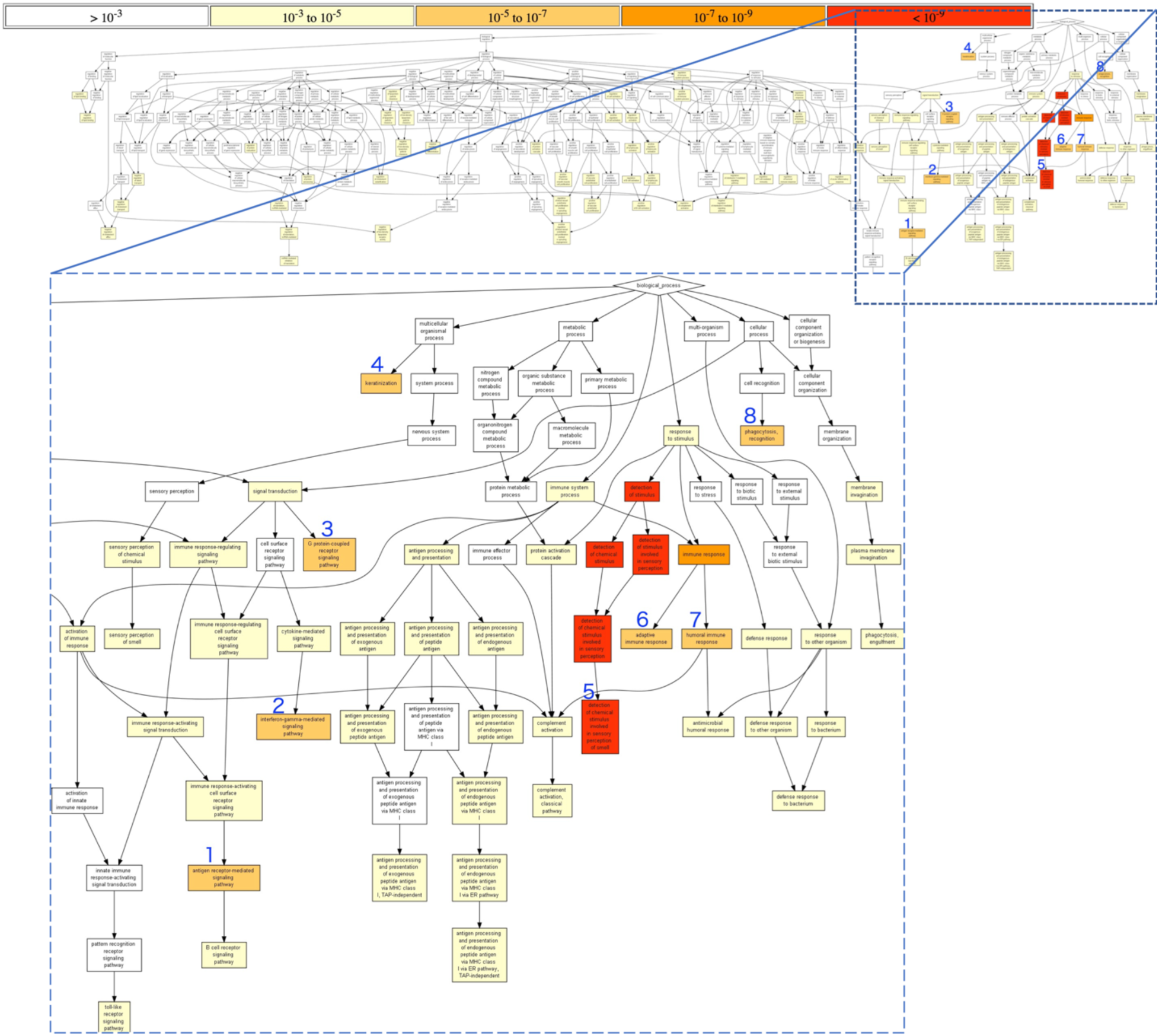
Biological process pathways with p values < 10^-5^. Regulation of Biological Processe (left branch) omitted due to p values > 10^-5^. Blue numbers = Gene ontology paths. Path 1 = Cellular process → signal transduction → immune response-regulating signaling pathway → immune response-regulating cell surface receptor signaling pathway → immune response-activating cell surface receptor signaling pathway → antigen receptor-mediated signaling pathway. Path 2 = Cellular process → signal transduction → cell surface receptor signaling pathway → interferon-gamma-mediated signaling pathway. Path 3 = Cellular process → signal transduction → G protein-coupled receptor signaling pathway. Path 4 = Multicellular organismal process → keratinization. Path 5 = *Response to stimulus → detection of stimulus → detection of chemical stimulus | detection of stimulus involved in sensory perception → detection of chemical stimulus involved in sensory perception → detection of chemical stimulus involved in sensory perception of smell*. Path 6 = Response to stimulus → immune response → adaptive immune response. Path 7 = Response to stimulus → immune response → humoral immune response. Path 8 = Cellular process → cell recognition → phagocytosis, recognition. Red font = p < 10^-9^

**Table 2.** Gene ontology terms with description, *p*-value, false discovery rate (FDR) *q*-value, and enrichment metrics.

| GO term | Description | $P$ Value | FDR $q$ Value | Enrichment (N, B, n, b)* |
| --- | --- | --- | --- | --- |
| GO:0050911 | detection of chemical stimulus involved in sensory perception of smell | 9.24E-14 | 5.87E-10 | 3.35<br>(1055,29,304,28) |
| GO:0050907 | detection of chemical stimulus involved in sensory perception | 7.04E-13 | 2.24E-09 | 11.34<br>(1055,31,45,15) |
| GO:0050906 | detection of stimulus involved in sensory perception | 7.60E-12 | 1.61E-08 | 10.05<br>(1055,35,45,15) |
| GO:0009593 | detection of chemical stimulus | 1.29E-11 | 2.06E-08 | 9.77<br>(1055,36,45,15) |
| GO:0051606 | detection of stimulus | 2.20E-10 | 2.80E-07 | 7.66<br>(1055,49,45,16) |
| GO:0006955 | immune response | 8.05E-08 | 8.53E-05 | 9.77<br>(1055,72,15,10) |
| GO:0002250 | adaptive immune response | 1.04E-07 | 9.42E-05 | 7.54<br>(1055,30,56,12) |
| GO:0007186 | G protein-coupled receptor signaling<br>pathway | 4.11E-07 | 3.27E-04 | 4.94<br>(1055,76,45,16) |
| GO:0006959 | humoral immune response | 6.90E-07 | 4.87E-04 | 34.25<br>(1055,14,11,5) |
| GO:0050851 | antigen receptor-mediated signaling<br>pathway | 7.16E-07 | 4.55E-04 | 22.21<br>(1055,19,15,6) |
| GO:0006910 | phagocytosis, recognition | 2.71E-06 | 1.57E-03 | 87.92 (1055,4,9,3) |
| GO:0031424 | keratinization | 6.85E-06 | 3.63E-03 | 11.81 (1055,8,67,6) |
| GO:0060333 | interferon-gamma-mediated signaling<br>pathway | 8.47E-06 | 4.14E-03 | 35.17 (1055,8,15,4) |
| GO:0051249 | regulation of lymphocyte activation | 1.01E-05 | 4.57E-03 | 15.63<br>(1055,27,15,6) |
| GO:0006958 | complement activation, classical pathway | 1.18E-05 | 4.99E-03 | 70.33 (1055,5,9,3) |
| GO:0007606 | sensory perception of chemical stimulus | 1.27E-05 | 5.04E-03 | 16.91 (1055,8,39,5) |
| GO:0002429 | immune response-activating cell surface receptor signaling pathway | 1.29E-05 | 4.83E-03 | 15.07<br>(1055,28,15,6) |
| GO:0002768 | immune response-regulating cell surface receptor signaling pathway | 2.51E-05 | 8.88E-03 | 13.61<br>(1055,31,15,6) |
| GO:0002694 | regulation of leukocyte activation | 2.51E-05 | 8.41E-03 | 13.61<br>(1055,31,15,6) |
| GO:0042742 | defense response to bacterium | 2.98E-05 | 9.46E-03 | 31.26 (1055,15,9,4) |

\*N = Total number of genes; B = Number of genes in the total number of genes associated with a particular GO term; n = number of these genes in the target set; b = number of genes from the target set associated with the GO term

As the Joint Pathway Analysis in MetaboAnalyst does not rank genes, a list of combined genes and metabolites obtained with the output from the exome and metabolomic analyses was filtered for a ranked score cutoff of 1.0, arriving at 260 genes and the eight relevant metabolites, which are pyridoxine, kynurenic acid, acetylcysteine, citraconic acid, phosphocreatine, hippuric acid, xylitol, and 2-deoxyuridine. This list was entered into MetaboAnalyst. A Holm *p* value of 0.05 was set as a cut-off for the enriched metabolite sets. Of the resulting top ten enriched metabolite sets, one was olfactory transduction, seven were directly related to the immune or autoimmune system (graft-versus-host disease, allograft rejection, antigen processing, type 1 diabetes, autoimmune thyroid disease, asthma, phagosome), or parasitic (Leishmaniasis) infection. Table 3 lists the GO terms obtained with these parameters.

**Table 3.**
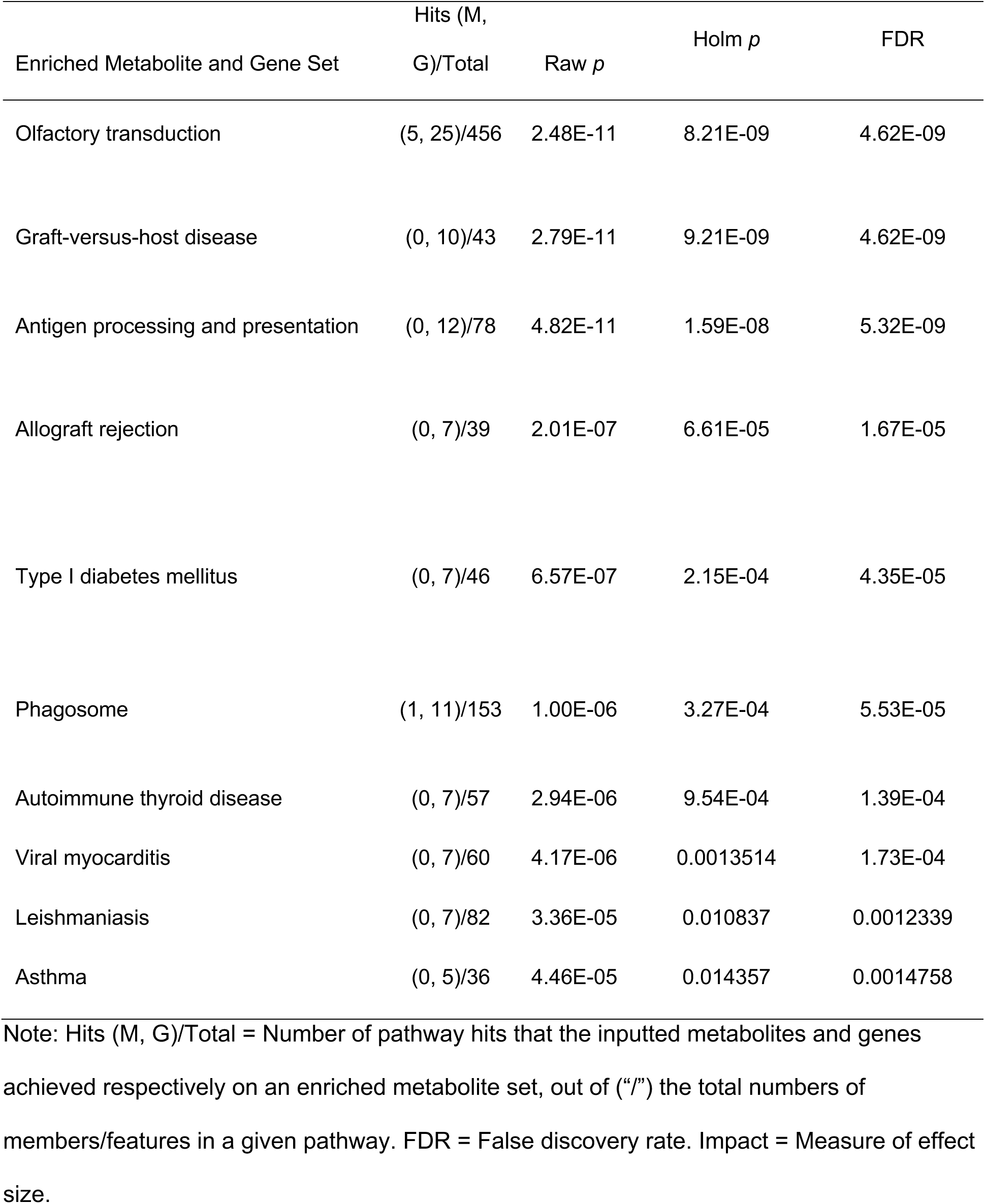
Output of combined exome and metabolome gene ontology analysis.

## DISCUSSION

In this pilot study, exome and metabolomic case/control analyses were applied individually and jointly to samples from participants with and without dyslexia. To our knowledge, this represents the first study of its kind in the field of dyslexia.

### Combined Metabolite and Exome Analysis

Preliminary results emerged regarding pathways at the intersection of the genomics and metabolomics analysis that are potentially novel. The most significant finding in the joint analysis was the olfactory transduction pathway. This pathway was linked with both genes and metabolites, with the input metabolites associated with five genes in the olfactory transduction pathway, and the input genes overlapping with 25 genes in the olfactory transduction pathway. Genes identified in the joint pathway analysis primarily include those in the OR (olfactory receptor) 1, 5, and 8 subfamilies. Although this pathway has not been previously associated with dyslexia, it has been implicated in other neurological disorders such as ADHD, ASD, and obsessive-compulsive disorder (OCD) (Crow et al., 2020; Karsz et al., 2008). Another study, using an olfactory ERP design, reported substantial evidence for the alteration of the later (more than 542 ms past stimulus onset), modality-general processing stage of olfactory processing in individuals with ASD (Okumura et al., 2020). Although there is no evidence to date supporting the link between the olfactory sensory pathway and dyslexia, there is evidence supporting the link between general sensory processing pathways and dyslexia. As mentioned, an fMRI study of auditory and visual stimuli showed that adults with dyslexia responded with significantly less neural adaptation to repetitive stimuli, compared to typical adults, regardless of type of stimulus, for example, written words, pictures of faces (Perrachione et al., 2016), and an ERP study showed diminished auditory gating in adults with dyslexia (Peter, McCollum, et al., 2019). Also note the relative abundance of sensory dysregulation and difficulties with attention and/or focus in the participants with dyslexia in the present study (Appendix). Given this context, the present study may be the first to suggest a link between dyslexia and the olfactory transduction pathway.

The other seven pathways were either directly or indirectly associated with the immune or autoimmune systems. Note that one of the participants in the dyslexic group (see Appendix) had Hashimoto’s disease, a severe form of hypothyroidism. A recent study found a high prevalence of thyroid autoimmunity disorders among individuals with dyslexia (Degrandi et al., 2021).

Several studies have found links between the olfactory sensory pathway and hypothyroidism. In one such study, there was a statistically significant difference in the recorded cortical responses between groups of patients with overt clinical hypothyroidism versus subclinical hypothyroidism. The authors state that “registration of cortex potentials at irritation of olfactory [nerves] offers possibilities for using this method as an objective indicator of hypothyroidism severity and prognostic process factor” (Świdziński et al., 2016). In the overt hypothyroidism groups, they found longer latency times of olfactory potentials PN1 and PN5 that were statistically significant to both the subclinical hypothyroidism and reference groups, which suggests that those with severe cases of hypothyroidism cannot appropriately respond to smell. Older studies have also established a link between hypothyroid mice and humans and malfunctioning olfactory responses (Świdziński et al., 2016).

Another pathway of significance is the Toxoplasmosis pathway, with our inputted metabolites associated with one gene in the Toxoplasmosis pathway, and our inputted genes overlapping with eight genes in the Toxoplasmosis pathway. Studies have shown that toxoplasmosis plays an important role in neurodevelopmental disorders such as schizophrenia and ASD (Al Malki et al., 2021; Fuglewicz et al., 2017; Nayeri et al., 2020; Torrey & Yolken, 2003). Although the relationship between prenatal toxoplasmosis and autism is unclear (Al Malki et al., 2021), there is a notable comorbidity between ASD and toxoplasmosis (Nayeri et al., 2020). Evidence also suggests a correlation between toxoplasmosis and schizophrenia, with negative effects on brain growth (Fuglewicz et al., 2017), and other studies have shown that an exposure to *Toxoplasma gondii* can affect behavior and neurotransmitter functions (Torrey & Yolken, 2003). Similar to the olfactory pathway, there is no substantial evidence to support significant involvement of the toxoplasmosis pathway in individuals with dyslexia, but evidence that supports a role for the toxoplasmosis pathway in neurodevelopmental disorders suggests further research is needed.

### Additional Findings from the Metabolome Analysis

Genes in the immune response and sensory perception pathways were implicated by the results of the metabolomic analysis. In addition, eight metabolites (kynurenic acid, pyridoxine, acetylcysteine, citraconic acid, 2-deoxyuridine, hippuric acid, phosphocreatine, and xylitol) were identified as markers for enhanced expression of dyslexia, as shown in Figures 1 and 2. The following is an exploration of possible relationship between these metabolites and dyslexia or other developmental disorders.

Findings in an animal study suggested that a significant increase in the expression of kynurenic acid, similar to the higher concentration of this metabolite in the participants with dyslexia in our study, may contribute to increased cognitive impairments, decreased performance in sustained attention tasks, and disrupted auditory gating (Kozak et al., 2014). A study of individuals with schizophrenia showed that an increase in kynurenic acid can disrupt prepulse inhibition, which is a behavioral model that measures sensorimotor gating (Erhardt et al., 2004). A study that measured kynurenic acid levels in school children with ASD, compared to a control group, found that the ASD group had significantly higher levels of kynurenic acid than the control group, consistent with the present findings in the dyslexia group (Bilgiç et al., 2020). Although no prior literature was found regarding any association between dyslexia and kynurenic acid, we infer that kynurenic acid could play a significant role in the development of dyslexia, as our phenotypic data resemble symptoms exhibited by schizophrenic patients, especially with respect to attention deficit and motor problems.

Additionally, although not specifically identified or associated with dyslexia, some studies have found abnormally high levels of pyridoxine (Vitamin B6) in individuals with ASD. According to one study, these results are consistent with the hypothesis that pyridoxal kinase, which regulates the vitamin B6 metabolism and is involved in the regulation of over 100 neurotransmitters, is severely inhibited in individuals with ASD (Adams et al., 2006). The small size of our study preludes the conclusion that pyridoxal kinase is implicated in the development of dyslexia; however, future research should investigate the role of pyridoxal kinase in dyslexic patients, as pyridoxal kinase and the vitamin B6 metabolism are associated with ASD, schizophrenia, Alzheimer’s disease, and Parkinson’s disease (Mascolo et al., 2019).

Acetylcysteine has been demonstrated to ameliorate autism symptoms in children. Six to eight weeks of N-acetylcysteine (NAC) supplementation has been shown to improve social awareness, irritability, and hyperactivity in children with ASD (Lee et al., 2021). Acetylcysteine has also been used to treat individuals with OCD, and results showed that NAC was well-tolerated and effectively reduced symptom severity (Oliver et al., 2015). In our analysis, however, acetylcysteine levels were higher in the dyslexia group, compared to the control group. Nevertheless, the link between acetylcysteine and dyslexia is still unexplored, and further metabolite studies on acetylcysteine and its role in the development and pathogenesis of dyslexia are justified.

Phosphocreatine is another metabolite of interest in our study for which we were not able to. find literature characterizing its role in dyslexia. However, phosphocreatine has been found to play a role in ASD. This metabolite was found in lower concentrations children and adolescents with ASD, compared to typical controls (Golomb et al., 2014). This is in contrast with our findings, where higher concentrations of phosphocreatine were seen in the dyslexia group, compared to the control group. In another study, phosphocreatine was found in lower concentrations in white matter tracts in adolescents with ASD (O’Neill et al., 2020). This is another metabolite for which the link between phosphocreatine and dyslexia invites further exploration.

Hippuric acid has also been implicated in ASD. In two studies, urinary concentrations of 3-Hydroxyhippuric Acid were found in higher concentrations in children with ASD, compared to typical peers (Lis et al., 1976; Xiong et al., 2016). In the present study, hippuric acid was found in higher concentrations in the dyslexia group, compared to the control group, which is consistent with the ASD study findings. Therefore, it is reasonable to investigate this metabolite in larger dyslexia cohorts.

Although citraconic acid has not been studied extensively in neurodevelopmental disorders, we did find that it interacts with other metabolites in the citric acid cycle that were identified in our analysis, including deoxyuridine, D-Xylitol, acetylcysteine, and pyridoxine. Citric acid intermediates and products such as pyruvic acid, NADP, oxygen, adenosine diphosphate (ADP), adenosine triphosphate (ATP), coenzyme A (CoA), and NADH are all associated in metabolite-metabolite interactions of our selected metabolites. Although the citric acid cycle has not been previously implicated in the pathogenesis of dyslexia, this evidence suggests that the Krebs cycle may play a role in it.

In children with ASD, carboxycitric acid, which is a product of the Krebs cycle, has been identified as a potential biomarker (Lis et al., 1976; Xiong et al., 2016). The Krebs cycle has also been associated with mitochondrial disorder in individuals with ASD, where urinary TCA cycle intermediates were found elevated in those with autism (Griffiths & Levy, 2017; Lombard, 1998). TCA cycle intermediates such as 2-oxoglutarate, cis-aconitate, and succinate, along with glutamate, acetate, and lactate, were found at increased levels in children with ASD compared to typical controls (Orozco et al., 2019).

Our metabolite-metabolite analysis was performed to identify intermediates and shared pathways. Many of the intermediates identified in our analysis are related to or directly interact within the Krebs (or TCA) cycle, illustrating the TCA cycle’s potential role in the pathogenesis of dyslexia. Notably, we identified citraconic acid, pyridoxine, xylitol, acetylcysteine, and deoxyuridine in our analysis as metabolites of interest, as they are all involved with Krebs cycle intermediates.

Glutamate, a product of the Krebs Cycle, has been specifically implicated in dyslexia (Del Tufo et al., 2018; Hancock et al., 2017; Pugh et al., 2014). The findings in the present study, however, did not implicate glutamate. Further research is needed to resolve this discrepancy and to characterize the role of glutamate is a key driver in the pathogenesis of dyslexia.

### Additional Findings from the Exome Analysis

In the exome analysis, Gorilla identified several genes related to immune responses pathways, especially those influencing antigen processing and the adaptive immune response, in particular, the HLA gene family, especially HLA-B, HLA-C, HLA-DRB, and the HLA-DQ subfamilies. This family has previously been implicated in disorders of spoken and written language (Nudel et al., 2014). Parent-of-origin effects between HLA-B B8 and HLA-DQA1 and receptive language ability were found in individuals with disorders of spoken language, implicating specifically the DR10 allele of HLA-DRB1 (Nudel et al., 2014). Consequently, the DQ and DRB subfamilies of HLA, and possibly the B and C subfamilies as well, may play a major role in language disorders including dyslexia.

The exome analysis revealed additional pathways that are potentially associated with dyslexia. In particular, signal transduction and interferon-gamma-mediated signaling have been associated with various neurodevelopmental disorders. One such study found that IFN-γ “disproportionately altered the expression of genes associated with schizophrenia and autism,” and, therefore, was involved in interferon signaling in neurodevelopmental disorders (Warre-Cornish et al., 2020). Another study identified the dysregulation of many key signaling pathways, including WNT, BMP, SHH, RA signaling in individuals with ASD (All of Us Research Program et al., 2019; Kumar et al., 2019). We have shown in a previous study using evoked response potentials and an auditory gating paradigm that adults with dyslexia had deficits in neural adaptations similar to those found in adults with schizophrenia (Peter, McCollum, et al., 2019). This would suggest potential common pathways between multiple neurological disorders such as dyslexia and schizophrenia.

### Limitations, Conclusions, and Future Studies

The main limitation of this pilot study is the small sample size and the inherently reduced statistical power. In addition, whole exome analysis was conducted, not whole genome analysis (WGS). WGS could identify potentially rare or novel structural or noncoding variants that might contribute to the phenotype, but the burden on statistical power would be even greater. Further, some participants were biologically related, which reduces the independence of each observation. Specifically, both our metabolite analysis using MetaboAnalyst and exome analysis using PLINK/GOrilla did not account for the relatedness among some of the participants. The reduced independence of observations could potentially result in overcorrection for multiple testing through commonly used approaches, such as the Bonferroni correction, which has been found to be too conservative due to the presence of both linkage disequilibrium (LD) and related individuals (Wang, 2011). As a result, the corrected p-values reported here could be too conservative.

Another limitation of our study is with the nature of the metabolites – as metabolite levels were taken from saliva, certain metabolites are sensitive to environmental changes. As we could not control for what our participants consumed prior to our testing, certain compounds such as caffeine and sweeteners such as xylitol can be influenced by environmental factors (Gómez-Fernández et al., 2021). Since xylitol has been identified in our study as a metabolite of interest, it may not be possible for us to properly determine if this metabolite, along metabolites that can change drastically due to environmental factors, can be associated with dyslexia. In summary, this combined metabolomics and exome analysis pilot study found suggestive evidence for involvement of sensory dysregulation and immune and autoimmune systems in dyslexia that could be investigated in larger participant samples.

## Acknowledgments

Many thanks to the participants for their time and efforts. This work was supported by the following grants: Arizona State University JumpStart Fund to B. Peter and V. Dinu; University of Washington Royalty Research Fund to B. Peter; National Institutes of Health R01HD088431 to W. Raskind.

## Web Resources

Agilent MassHunter Software: https://www.agilent.com/en/promotions/masshunter-mass-spec

BioMart: http://www.biomart.org/

GOrilla: http://cbl-gorilla.cs.technion.ac.il/

MetaboAnalyst 5: https://www.metaboanalyst.ca/MetaboAnalyst/home.xhtml

PLINK: http://zzz.bwh.harvard.edu/plink/

R: https://www.r-project.org

R Studio: https://www.rstudio.com/

SeattleSeq Annotation Server: https://snp.gs.washington.edu/SeattleSeqAnnotation154/

## Data and Code Availability

The metabolome data are available in the supplemental information as Table S1. The code is available there as Document S1. The exome data have not been uploaded into a public database because they were not obtained with federal funding, but they are available from the corresponding author on request.

Appendix. Participants by family, codes, sex, dyslexia affectation, additional traits, age at exome collection and metabolome analysis, and available biological sample

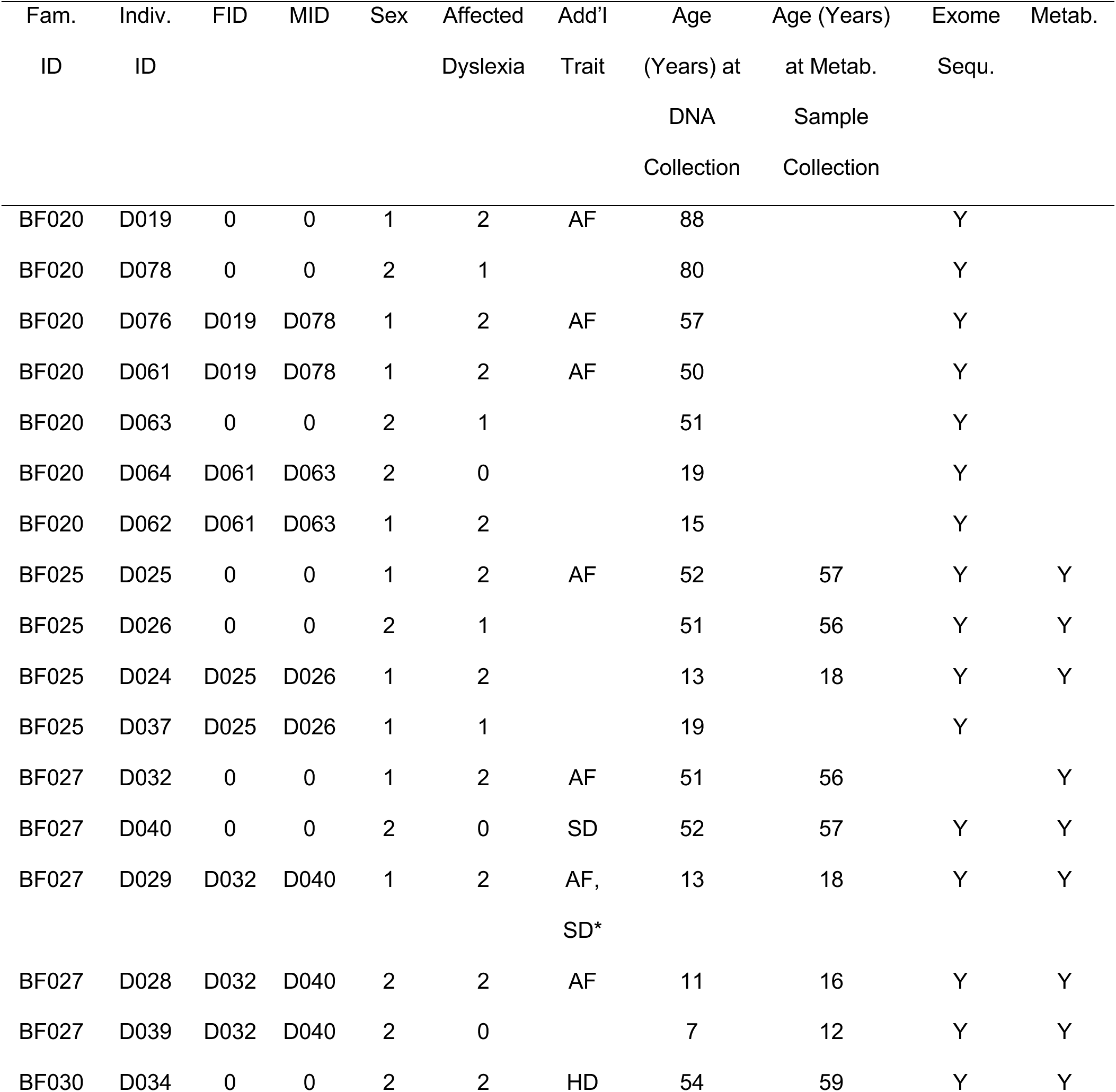
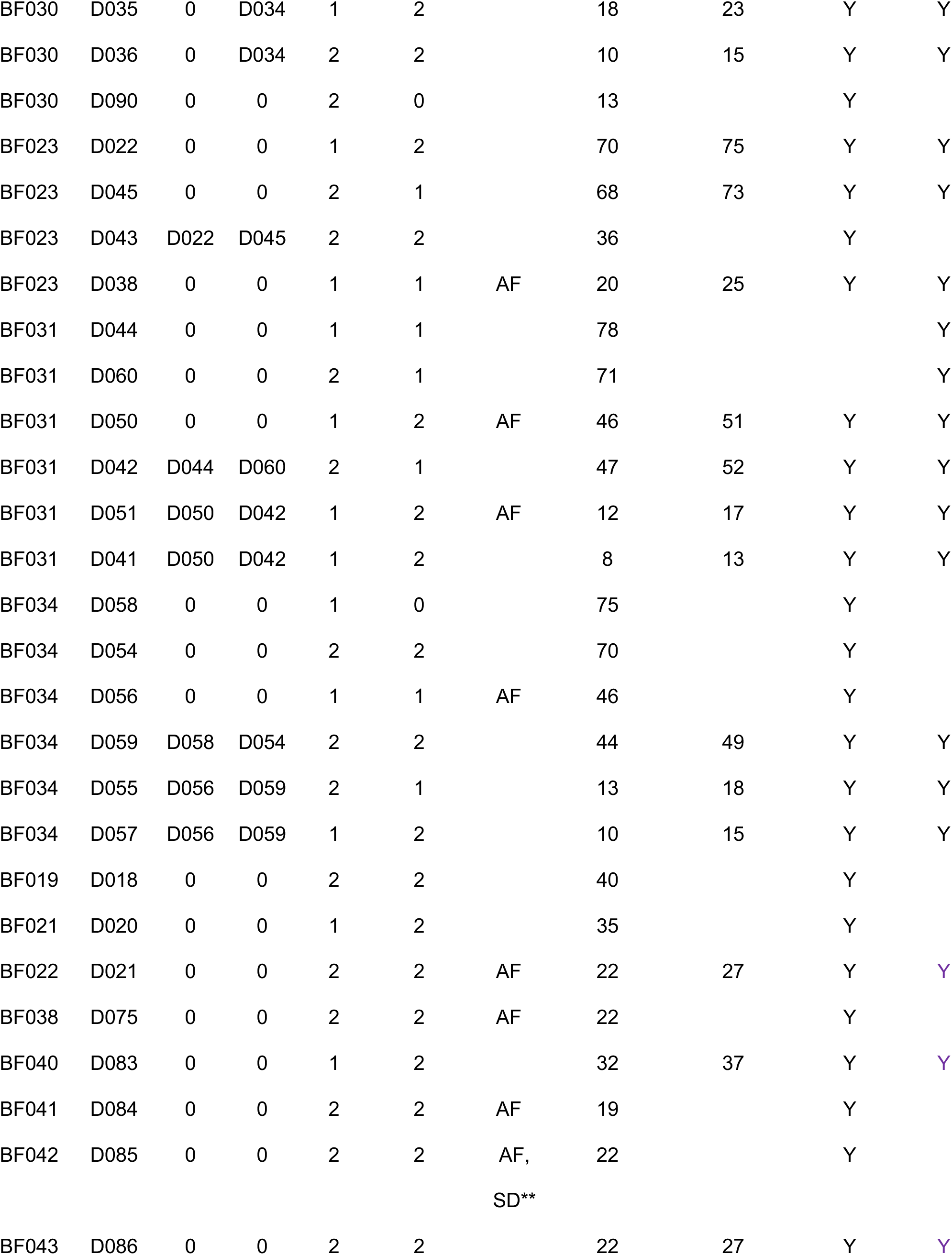
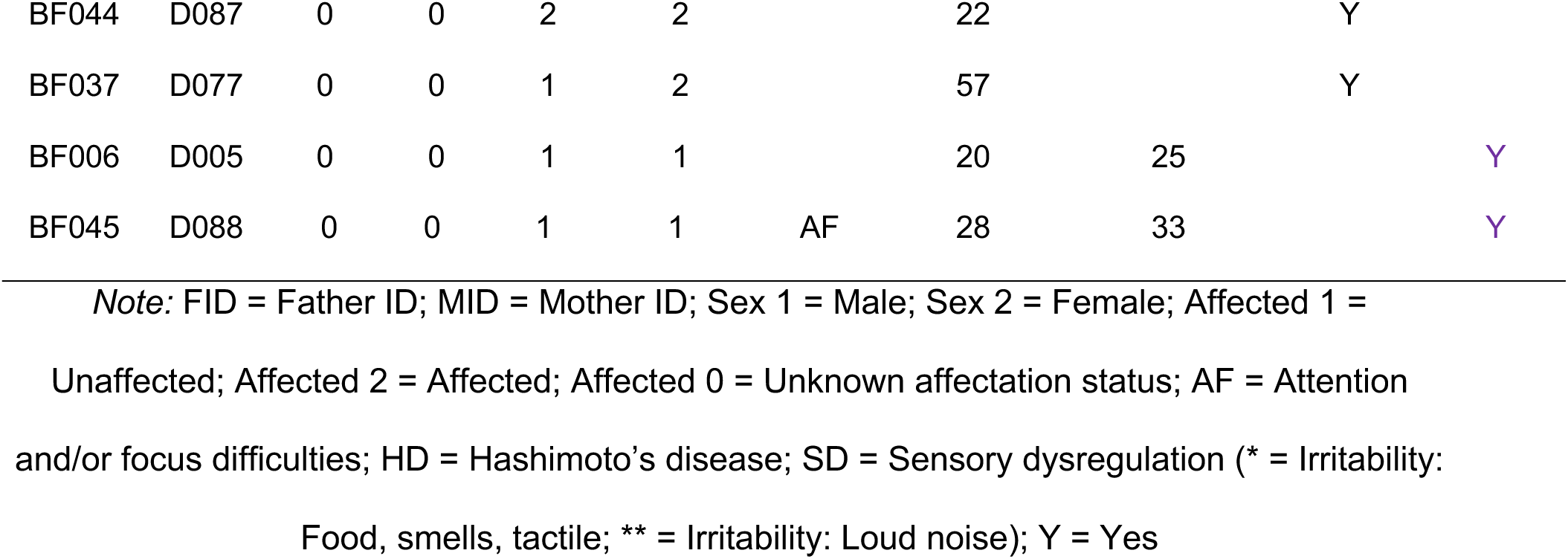

