## Supplementary material for "Joint exome and metabolome analysis in individuals with dyslexia: Evidence for associated dysregulations of olfactory perception and autoimmune functions": Document S1

### Libraries

library(biomaRt)

library(dplyr)

library(rlist)

library(rlang)

library(EnsDb.Hsapiens.v79)

library("openxlsx")

require(data.table)

library(tidyverse)

library(goseq)

library("org.Hs.eg.db")

library(EDASeq)

### Pick mart

ensembl <- useMart("ENSEMBL\_MART\_SNP", dataset = "hsapiens\_snp", host="grch37.ensembl.org")

ensembl

### ensembl <- useMart("ensembl", dataset = "hsapiens\_gene\_ensembl")

### Read as2.assoc file from pLink analysis

as2 <- read.table("final\_dyslexia\_results.assoc", sep="", header=TRUE)

#Only get SNPs w/ P-values < 0.05

snps <- dplyr::filter(as2, P < 0.05)

#unique(snps)

### First, get the genes associated with SNPs without RS labels

getNonRstoGenes <- function () {

  # Get only SNPs without an RS tag (these are signified as dots on the .assoc file)

  alldots <- snps %>% dplyr::filter(SNP == ".")

  # Set up parameters

  mart <- useMart("ensembl", host="grch37.ensembl.org")

  mart <- useDataset("hsapiens\_gene\_ensembl", mart)

  attributes <- c("ensembl\_gene\_id","start\_position","end\_position","strand","hgnc\_symbol","chromosome\_name")

  filters <- c("chromosome\_name","start","end")

  values <- data.frame(alldots$CHR, alldots$BP, alldots$BP + 1)

  all\_genes = data.frame(ensembl\_gene=character(), start=character(), end=character(), strand=character(), hgnc\_symbol=character(), chromosome\_number=character())

  print(lengths(all\_genes)[1])

  # loop thru RS IDs

  for (i in c(1:lengths(values)[1])) {

    print (i)

    all.genes <- getBM(attributes=attributes, filters=filters, values=list(values[i, ]$alldots.CHR, values[i, ]$alldots.BP, values[i, ]$alldots.BP + 1), mart=mart, useCache = FALSE)

    if (dim(all.genes)[1] != 0) {

      all.genes <- unique(all.genes)

      all\_genes[i, ] <- all.genes

      print(all.genes)

    }

  }

  all\_genes

  # Write SNP (non-RS) to genes (duplicates removed)

  write.xlsx(unique(all\_genes), file = "snps\_nonRS\_to\_genes.xlsx",

             sheetName = "by\_snp", append = FALSE)

}

getNonRstoGenes()

gRStoGenes <- function() {

  #Get only SNPs with RS

  allRS <- (snps$SNP)[snps$SNP %in% "." == FALSE]

  # Get gene IDs with SNP

  geneIds <- getBM(attributes=c(

    "refsnp\_id", "ensembl\_gene\_stable\_id"),

    filters="snp\_filter", values=allRS,

    mart=ensembl, uniqueRows=TRUE, useCache = FALSE)

  geneIDsA <- ensembldb::select(EnsDb.Hsapiens.v79, keys=geneIds$ensembl\_gene\_stable\_id, keytype = "GENEID", columns = c("SYMBOL","GENEID"))

  geneIDsA <- unique(geneIDsA)

  # Write SNP to genes (symbols) (duplicates removed)

  write.xlsx(unique(data.table(cbind(refsnp\_id = geneIds$refsnp\_id, gene\_symbol = geneIDsA$SYMBOL[match(geneIds$ensembl\_gene\_stable\_id, geneIDsA$GENEID)]), key="refsnp\_id")), file = "C:\\Users\\14129\\Junior Year\\Dyslexia\\Results\\Exomes Results\\Results from Exomes\\snps\_to\_genes\_SYMBOL.xlsx",

             sheetName = "by\_snp", append = FALSE)

  # Write Gene (ENSEMBL) to SNP

  write.xlsx(data.table(geneIds, key="ensembl\_gene\_stable\_id"), file = "C:\\Users\\14129\\Junior Year\\Dyslexia\\Results\\Exomes Results\\Results from Exomes\\genes\_to\_snps\_ENSEMBL.xlsx", 

             sheetName="by\_ensembl", append=FALSE)

  # Write genes (symbols) to snp (duplicates removed)

  write.xlsx(unique(data.table(cbind(refsnp\_id = geneIds$refsnp\_id, gene\_symbol = geneIDsA$SYMBOL[match(geneIds$ensembl\_gene\_stable\_id, geneIDsA$GENEID)]), key="gene\_symbol")), file = "C:\\Users\\14129\\Junior Year\\Dyslexia\\Results\\Exomes Results\\Results from Exomes\\genes\_to\_SNP\_SYMBOL.xlsx",

             sheetName = "by\_symbol", append = FALSE)

}

RStoGenes()

### To see all the top gene Id using SYMBOL

genestosnpsymbol <- openxlsx::read.xlsx("C:\\Users\\14129\\Junior Year\\Dyslexia\\Results\\Exomes Results\\Results from Exomes\\genes\_to\_SNP\_SYMBOL.xlsx")

genestononsnp <- openxlsx::read.xlsx("C:\\Users\\14129\\Junior Year\\Dyslexia\\Results\\Exomes Results\\Results from Exomes\\snps\_nonRS\_to\_genes.xlsx")

length(c(genestosnpsymbol$gene\_symbol, genestononsnp$hgnc\_symbol))

write.csv(sort(table(c(genestosnpsymbol$gene\_symbol, genestononsnp$hgnc\_symbol)), decreasing=TRUE), file = "C:\\Users\\14129\\Junior Year\\Dyslexia\\Results\\Exomes Results\\Results from Exomes\\FINAL\_table\_snp\_to\_genes\_SYMBOL.csv")

### combine both gene

sort(table(c(genestosnpsymbol$gene\_symbol, genestononsnp$hgnc\_symbol)), decreasing = TRUE)

combinedGeneSymbol <- sort(table(c(genestosnpsymbol$gene\_symbol, genestononsnp$hgnc\_symbol)), decreasing = TRUE)

length(combinedGeneSymbol)

lengths(genestononsnp)

lengths(genestosnpsymbol)

ensemblmart <- useDataset('hsapiens\_gene\_ensembl', mart)

listSymbolNames <- attributes(combinedGeneSymbol)

listSymbolNames

### use biomaRt to get gene length by symbol

symbolGeneLength <- getBM(attributes=c('hgnc\_symbol', 'chromosome\_name', 'start\_position', 'end\_position', 'strand'),

      filters=c('hgnc\_symbol'),

      values=unlist(listSymbolNames),

      mart=ensemblmart, useCache = FALSE)

### Gene size

symbolGeneLength$length <- symbolGeneLength$end\_position - symbolGeneLength$start\_position

symbolGeneLength$hgnc\_symbol

symbolGeneLength$chromosome\_name <- as.numeric(as.character(symbolGeneLength$chromosome\_name))

### Keep only unique gene symbols

symbolGeneLength <- symbolGeneLength[!duplicated(symbolGeneLength$hgnc\_symbol), ]

inputGeneList <- read.csv("FINAL\_table\_snp\_to\_genes\_SYMBOL.csv")

inputGeneList

#inputGeneList$length <- NA

colnames(inputGeneList) <- c("X", "hgnc\_symbol", "length")

finalList <- merge(x=inputGeneList, y=symbolGeneLength, by.y = "hgnc\_symbol", by.x = "hgnc\_symbol")

### Create ranked score function

finalList$rankedScore <- finalList$length.x/finalList$length.y\*10000

finalList

### order final list in decreasing order by rankedScore

finalList <- finalList[order(finalList$rankedScore, decreasing=TRUE), ]

write.csv(finalList, file = "rankedscores.csv")
